# Short- and long-term scaling behavior of blood pressure and pulse arrival time during sleep in healthy controls and patients with obstructive sleep apnea

**DOI:** 10.64898/2025.12.14.694239

**Authors:** Karsten Berg, Jan W. Kantelhardt, Martin Glos, Thomas Penzel, Niels Wessel, Ronny P. Bartsch

## Abstract

Obstructive sleep apnea (OSA) is characterized by recurrent respiratory events that trigger autonomic arousals and blood pressure (BP) surges, contributing to elevated cardiovascular risk. Photoplethysmography (PPG)-derived timing markers such as pulse arrival time (PAT) are frequently used as noninvasive surrogates of BP dynamics, yet their interpretation is confounded by the pre-ejection period and peripheral vascular effects. Here, we used detrended fluctuation analysis (DFA) to quantify short– and long-term scaling exponents of continuous blood pressure (Portapres), PPG-, and PAT-derived signals across sleep stages in healthy individuals and patients with OSA.

Directly measured systolic and diastolic BP exhibited a robust short– to long-term crossover across all sleep stages, with elevated short-range exponents (*α*_1_ *>* 1) and lower long-range exponents (*α*_2_ *<* 1), reflecting well-organized autonomic and vascular control. In OSA, this crossover persisted but was visibly attenuated, consistent with reduced short-term adaptability of cardiovascular regulation.

In contrast, PAT-based indices showed substantially weaker short-range correlations and minimal crossover structure. Systolic PAT displayed almost no separation between *α*_1_ and *α*_2_, and PPG-derived measures exhibited scaling patterns that differed fundamentally from BP. Across modalities, PAT (whether derived from BP or PPG) failed to reproduce the multiscale organization characteristic of beat-to-beat BP dynamics. Group comparisons further identified systolic BP scaling, particularly the short-range exponent *α*_1_, as the most sensitive marker of cardiovascular dysregulation in OSA, whereas PAT and PPG provided complementary but physiologically distinct information related to peripheral vascular and autonomic modulation.

These findings demonstrate that PAT and PPG timing measures should not be used as surrogates for BP in fractal or scaling analyses and underscore the unique diagnostic value of BP-derived scaling behavior for assessing cardiovascular regulation during sleep.

## 1 Introduction

Obstructive sleep apnea (OSA) is a severe sleep disorder characterised by repeated upper-airway collapses during sleep that lead to hypoxaemia, frequent arousals, and fragmented sleep architecture [1, 2]. It is highly prevalent, with recent estimates suggesting that nearly one billion people are affected globally [3]. Moreover, OSA is associated with hypertension, coronary disease, stroke, and increased mortality, making accurate detection and staging essential for clinical management [4]. Modern screening approaches increasingly rely on non-invasive physiological signals that are easier to deploy than full polysomnography (PSG) [5].

Fluctuations in blood pressure (BP) play a central role in the pathophysiology of OSA: repeated apnea events trigger sympathetic surges, transient BP elevations, and altered baroreflex sensitivity, linking OSA severity directly to cardiovascular risk [6, 7]. Therefore, continuous assessment of BP dynamics may provide complementary insight into autonomic cardiovascular regulation during sleep.

Photoplethysmography (PPG) has emerged as a promising tool for sleep monitoring and OSA detection [8]. Timing parameters derived from PPG and ECG, such as pulse arrival time (PAT) and pulse transit time (PTT), are particularly useful because they reflect aspects of cardiovascular regulation and autonomic control. PTT, defined as the time required for a pulse wave to travel between two arterial sites, inversely relates to arterial stiffness and BP [9]. PAT additionally includes the pre-ejection period (PEP) of the heart and therefore incorporates cardiac contractility and sympathetic activation [10, 11]. Since apnea episodes elicit characteristic autonomic and hemodynamic responses, both PAT and PTT exhibit measurable fluctuations during sleep-disordered breathing [12–14].

Beyond discrete event detection, the temporal organisation of cardiovascular signals provides additional insight into physiological regulation [15]. Detrended fluctuation analysis (DFA) quantifies the scaling of signal variability across different time ranges and yields short– and long-term scaling exponents (*α*_1_, *α*_2_) that relate to autonomic control mechanisms [16, 17]. Alterations in these exponents have been associated with impaired adaptability of cardiovascular and respiratory systems across sleep stages and disease states [18]. Recent developments in Network Physiology have expanded this perspective, showing how multiscale coupling, redundancy, and synergy shape cardiovascular dynamics under sleep and postural challenge [15, 19]. Additional work has demonstrated that physiological variability is best understood within a framework of constrained disorder, where deviations from healthy multiscale structure reflect impaired flexibility of autonomic and vascular control [20]. Together with advances in analyzing cardio-respiratory and vascular interaction gradients [19, 21], these insights highlight the need to evaluate cardiovascular signals not only through their average properties but through their multiscale organisation.

Following this multiscale/multi-organ approach, in a recent study, we examined DFA scaling behavior across several physiological systems, including beat-to-beat intervals, respiratory intervals, EEG amplitudes, and pulse transit times, and observed a consistent crossover from short-term to long-term correlations in most signals, except for PTT [16]. This absence of a clear *α*_1_-*α*_2_ crossover is in contrast with earlier findings for direct BP measurements and calls into question how well PAT/PTT serves as a surrogate for BP dynamics across time scales.

To address this, we analyzed data from the previous EU project DAPHNet [22], which uniquely combines full PSG and simultaneously recorded continuous non-invasive BP measurements. This configuration allows direct comparison of PAT and PPG-derived pulse dynamics with beat-to-beat BP fluctuations across sleep stages, and enables a systematic evaluation of whether PAT captures the multiscale temporal structure of BP in both healthy subjects and individuals with OSA.

## 2 Materials and Methods

### Dataset and Study Protocol

The study protocol and measurement setup of the DAPHNet project (2006-2009) have been described in detail elsewhere [22]. In brief, the study was approved by the ethics committee of the Charité-Universitätsmedizin Berlin, and all participants provided written informed consent. The original study investigated autonomic and cardiovascular responses to arousals in obstructive sleep apnea (OSA), using healthy subjects as controls. For the secondary analysis performed in this study, anonymized recordings were accessed in July 2022 for research purposes. The authors had no access to personal identifiers at any point during data handling or analysis.

Participants underwent two consecutive nights of full in-lab PSG (Embla Inc., Broomfield, CO). Recordings included EEG (C3/A2, C4/A1), bilateral EOG, chin and tibialis EMG, ECG, airflow, respiratory effort, SpO_2_, and finger photoplethysmography (PPG). Continuous non-invasive beat-to-beat blood pressure (BP) was recorded using a Portapres Model 2 device (Finapres Medical Systems), which alternates finger cuffs and applies hydrostatic height correction based on the volume-clamp method [23]. Night 1 served as adaptation; night 2 was manually scored according to AASM criteria. Of the original 44 subjects, 40 recordings were available for re-analysis: two were unavailable and two OSA subjects were excluded due to missing BP signals. Group-level descriptive statistics are provided in Table 1.

**Table 1.**
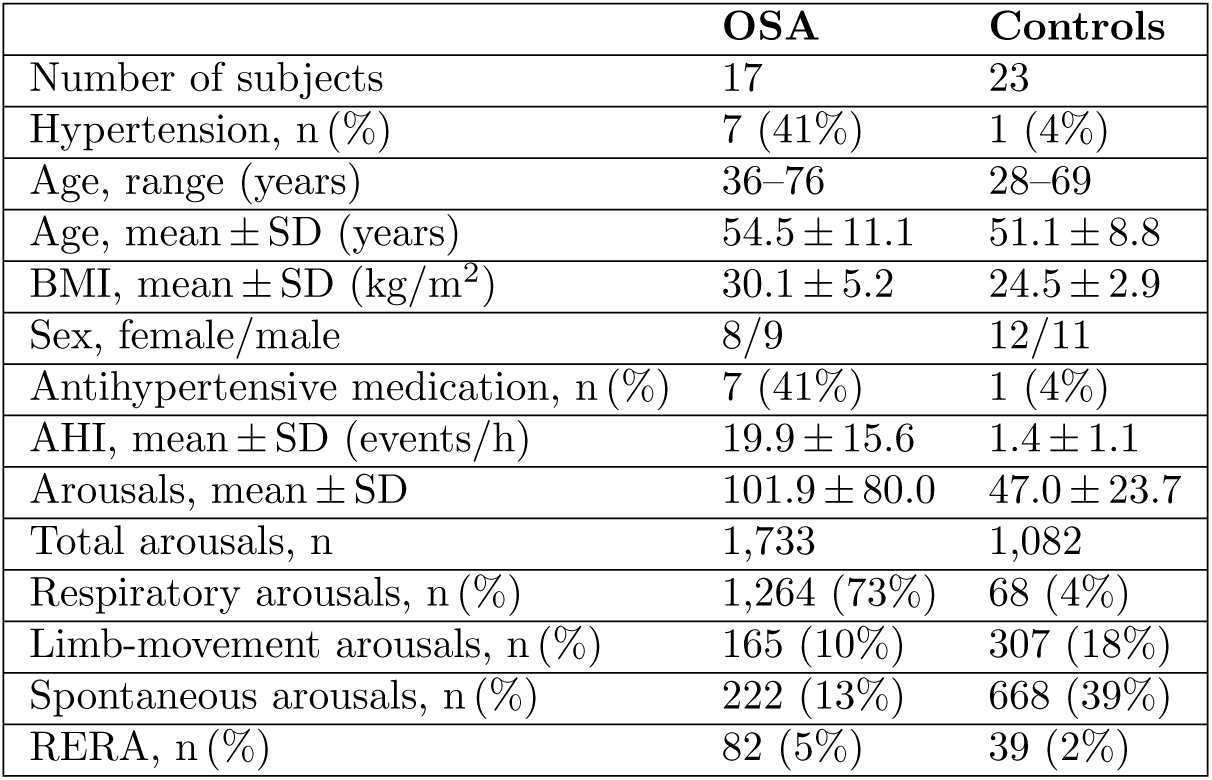
Demographic and clinical characteristics of OSA and control groups. BMI-Body Mass Index; AHI-Apnea–Hypopnea Index; RERA-Respiratory Effort–Related Arousal.

|  | OSA | Controls |
| --- | --- | --- |
| Number of subjects | 17 | 23 |
| Hypertension, n (%) | 7 (41%) | 1 (4%) |
| Age, range (years) | 36–76 | 28–69 |
| Age, mean ± SD (years) | 54.5 ± 11.1 | 51.1 ± 8.8 |
| BMI, mean ± SD (kg/m<sup>2</sup>) | 30.1 ± 5.2 | 24.5 ± 2.9 |
| Sex, female/male | 8/9 | 12/11 |
| Antihypertensive medication, n (%) | 7 (41%) | 1 (4%) |
| AHI, mean ± SD (events/h) | 19.9 ± 15.6 | 1.4 ± 1.1 |
| Arousals, mean ± SD | 101.9 ± 80.0 | 47.0 ± 23.7 |
| Total arousals, n | 1,733 | 1,082 |
| Respiratory arousals, n (%) | 1,264 (73%) | 68 (4%) |
| Limb-movement arousals, n (%) | 165 (10%) | 307 (18%) |
| Spontaneous arousals, n (%) | 222 (13%) | 668 (39%) |
| RERA, n (%) | 82 (5%) | 39 (2%) |

### Subjects

Controls were age– and sex-matched to OSA subjects. OSA was defined by an apnea–hypopnea index (AHI) *>*5 events/h (controls: AHI *<*5 events/h). Exclusion criteria followed [22] and included age *<*18 years, BMI *>*40 kg/m^2^, substance abuse, thyroid dysfunction, chronic pain, neurological or psychiatric disorders, significant cardiovascular or pulmonary disease, periodic limb movement index (PLMI) *>*5 events/h, and recent clinical trial participation or intercontinental travel.

### Physiological Signals

We analysed the following signals from night 2:

- ECG (200 Hz),
- PPG via pulse oximetry (100 Hz),
- continuous beat-to-beat BP from the Portapres system (200 Hz).

Signals were synchronised using internal device timestamps, compensating for known Embla system delays. Preprocessing steps included baseline removal, band-pass filtering and artefact suppression (full filter specifications in Supplementary Material).

### Morphologic Features

Figure 1 illustrates systolic and diastolic features of the BP pulse (as observed at the finger tip) and their relation to PPG morphology. Systolic BP (SBP) was defined as the pulse maximum; diastolic BP (DBP) as the subsequent local minimum on the falling limb.

**Fig 1.**
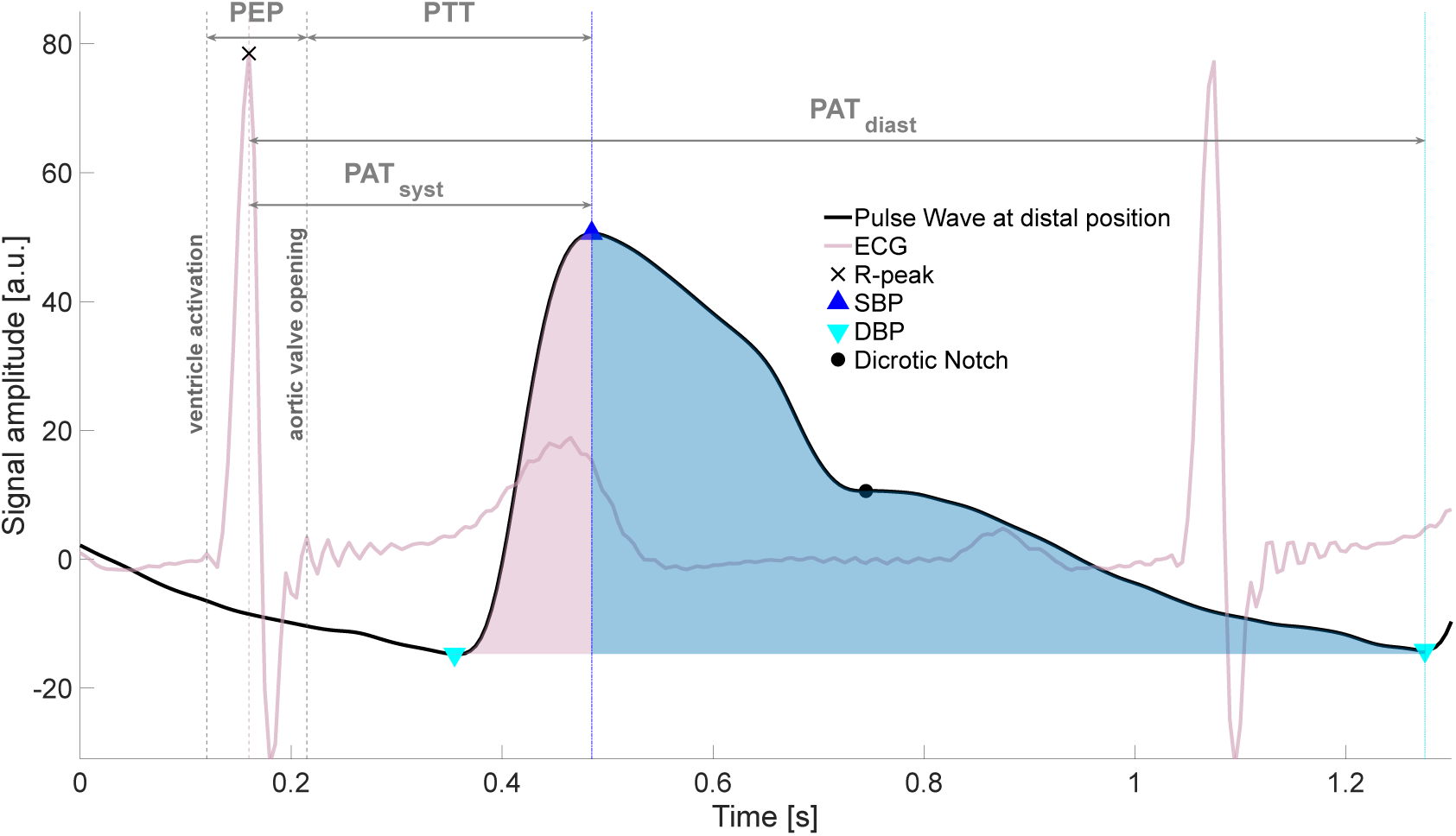
Characteristic features of the arterial blood pressure pulse wave measured at the finger tip. The red and blue shaded regions represent the anacrotic (rising) and catacrotic (falling) phases of the pulse wave, extending from the diastolic minimum (DBP) to the systolic maximum (SBP) and back. The first local minimum on the falling limb, the dicrotic notch, reflects aortic valve closure and marks the transition between systolic and diastolic components of the waveform. Pulse arrival time (PAT) is defined as the interval from a fiducial point within the QRS complex (typically the R-peak) to the arrival of the peripheral pulse waveform at its systolic or diastolic feature. PAT includes the pre-ejection period (PEP) and thus differs from pulse transit time (PTT), which reflects only the vascular propagation delay and requires identification of aortic valve opening or two peripheral PPG sensors for accurate estimation [10, 11]. Fiducial points are aligned with waveform peaks in this schematic for illustrative clarity. In our analysis, PAT_sys_ was defined as the interval from the R-peak to the SBP maximum in BP signals and to the systolic peak in PPG signals. PAT_dia_ was defined as the interval from the R-peak to the diastolic minimum (DBP in BP signals; first diastolic minimum following the systolic peak in PPG signals). The dicrotic notch was not used as a fiducial marker.

Pulse arrival times (PAT) were defined as:

- **PAT_sys_**: ECG R-peak to SBP (Portapres),
- **PAT_dia_**: ECG R-peak to DBP (Portapres).

The dicrotic notch was not used as a fiducial point. As PAT includes the pre-ejection period (PEP), it is physiologically distinct from pulse transit time (PTT), which requires two arterial measurement sites and reflects purely vascular propagation delay [10, 11]. For PPG signals, PAT_sys_ and PAT_dia_ were defined analogously as the intervals from the R-peak to the PPG systolic peak and to the first PPG minimum following that peak, respectively.

### Data Preprocessing

All analyses were performed in MATLAB (R2023a). ECG signals were corrected for baseline drift using cascaded median filters, and R-peaks were detected using the Benitez algorithm [24]. PPG and BP signals were filtered with zero-phase Butterworth filters to suppress high-frequency noise while preserving pulse morphology. Detailed filter orders, cutoffs, and resampling steps are provided in the Supplementary Material.

### Beat Segmentation and Validation

Beats were segmented using consecutive ECG R-peaks as boundaries. BP and PPG beats were aligned to the same R-peaks to ensure modality synchronisation. Beat morphology was assessed by correlating each beat with a subject-specific template; beats failing modality-specific correlation thresholds were excluded. SBP and DBP were identified using peak-valley detection refined within a ±0.2 s window around template extrema. Beats with physiologically implausible values (e.g. SBP *>*200 mmHg, DBP *<*10 mmHg) were discarded. RRIs, PATs, and BP values violating established physiological limits [16, 25] were excluded. Segments with missing signal or with calibration artifacts were also ignored (Fig. 2). Only uninterrupted sequences of ≥ 20 validated beats were used for further analysis.

**Fig 2.**
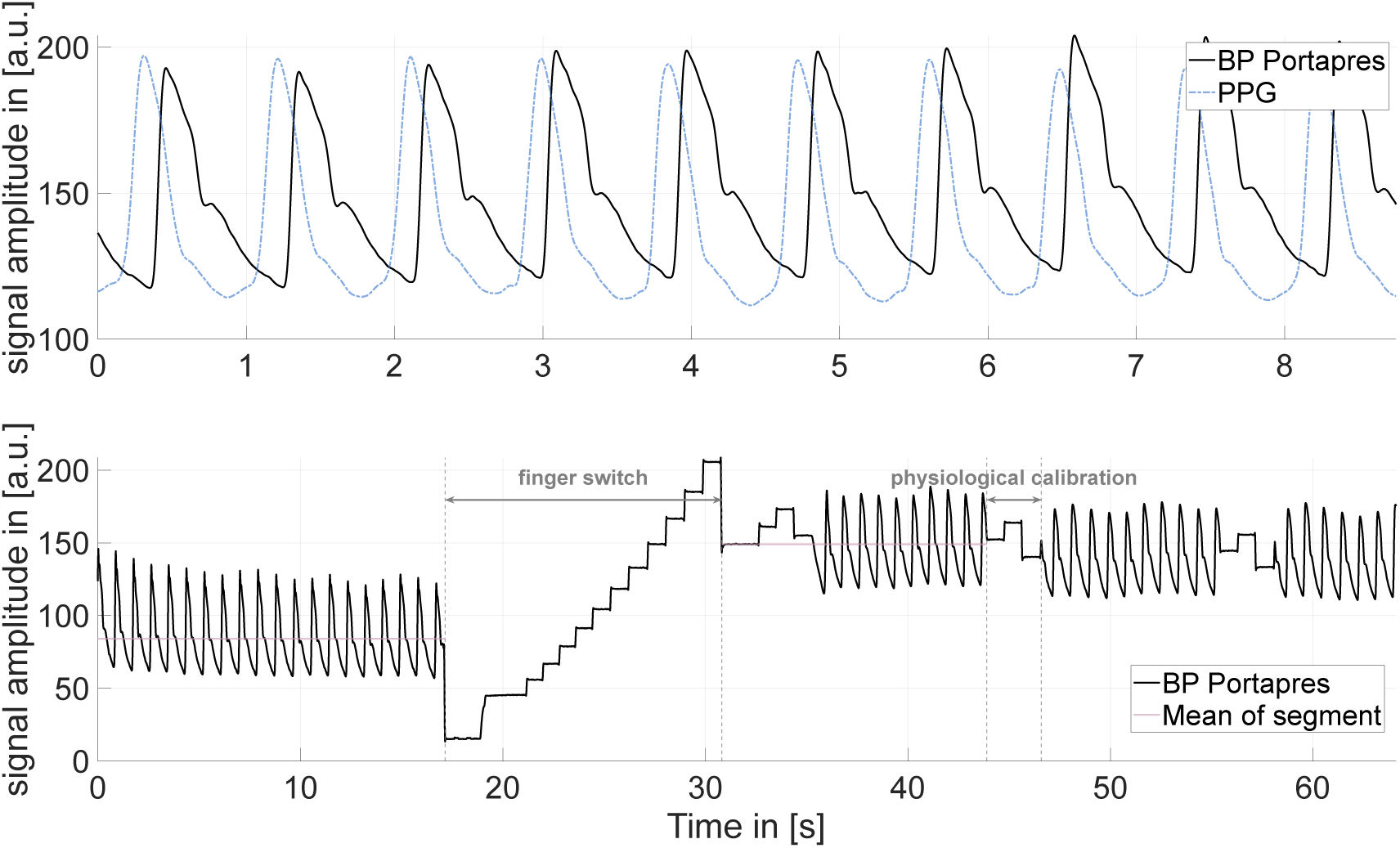
Typical artifacts in Portapres (BP) and PPG (pulse oximetry) recordings. *Top:* Example of simultaneous BP (solid line) and PPG (blue dashed line) waveforms. The PPG pulse wave lacks a distinct dicrotic notch and shows a temporal delay relative to the BP waveform. This absence of the dicrotic feature is likely attributable to age-related vascular stiffening, which attenuates the reflected wave component and highlights the physiological and morphological differences between the two signals. Whereas BP primarily reflects cardiac function and central arterial dynamics, the PPG pulse wave is more strongly influenced by peripheral vascular properties. *Bottom:* Representative artifacts in the Portapres BP signal. The device alternates measurement fingers approximately every 30 minutes to prevent vasoconstriction, producing abrupt baseline offsets. In addition, internal physiological recalibrations occur every 10–70 beats depending on signal quality [26, 27]. Both finger-switching and calibration events, as well as brief signal losses, lead to missing or invalid beats. In the present analysis, such segments were excluded, and only relative beat-to-beat changes in systolic and diastolic pressure (SBP and DBP) were evaluated, as these contain physiologically meaningful variability [28].

### Sleep Staging

Sleep stages were scored manually according to AASM rules. Because N1 episodes were sparse and typically too short for DFA, analyses focused on wake, N2, N3, and REM sleep.

### Detrended Fluctuation Analysis

Detrended fluctuation analysis (DFA) [29, 30] quantifies correlation properties of nonstationary time series. DFA2 (quadratic detrending) was applied to nine beat-to-beat time series: ECG-derived RRI; Portapres-derived SBP, DBP, PAT_sys_, PAT_dia_; and PPG-derived systolic peak, diastolic minimum, PAT_sys_, and PAT_dia_. Short-and long-term scaling exponents were defined as:

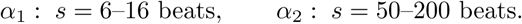

Only scaling fits with *r*^2^ *>* 0.9 were accepted (Fig. 3).

**Fig 3.**
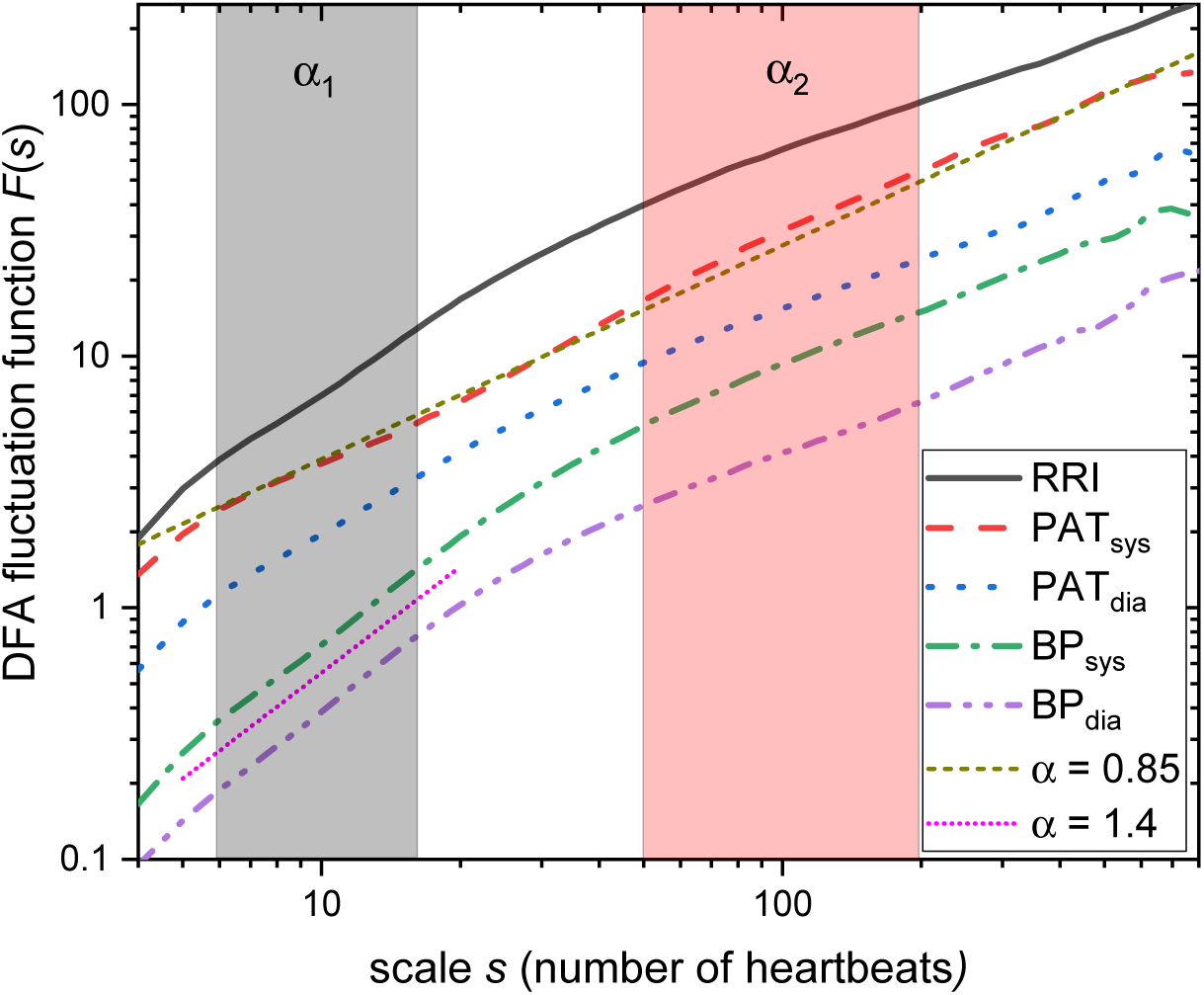
DFA fluctuation functions for cardiovascular time series. Double-logarithmic plots of the average DFA fluctuation function *F* (*s*) versus time scale *s* for 23 healthy subjects. Shown are the results for ECG-derived RR intervals (RRI), systolic and diastolic BP values from the Portapres device, and the corresponding systolic and diastolic pulse arrival times (PAT_sys_, PAT_dia_). BP-derived signals display strong short-term correlations (typical *α*_1_ ≈ 1.4) for both systolic and diastolic values, similar to the behavior observed in RRI. In contrast, PAT signals exhibit markedly weaker short-term correlations, and the crossover between short– and long-term scaling regimes is substantially reduced or absent, particularly for systolic PAT. Grey and red shaded regions indicate the fitting windows used to estimate the short-term (*α*_1_, 6–16 beats) and long-term (*α*_2_, 50–200 beats) scaling exponents, respectively. Straight reference lines with slopes *α* = 0.85 and *α* = 1.4 are shown as guides to the eye.

### Statistical Analysis

Because *α*_1_ and *α*_2_ were not normally distributed (Shapiro-Wilk, *p <* 0.05), non-parametric methods were used. The primary analyses focused on differences in scaling behavior across the physiological measures. Within each subject group (healthy, OSA), Kruskal-Wallis tests were applied across the five Portapres-derived measures (RRI, SBP, DBP, PAT_sys_, PAT_dia_) and across the four PPG-derived measures (systolic peak, diastolic minimum, PAT_sys_, PAT_dia_) for each sleep stage. Significant effects were examined using Bonferroni-corrected pairwise Wilcoxon tests. Stage-related differences within each physiological measure were assessed using Kruskal-Wallis tests across wake, REM sleep, N2, and N3, followed by Bonferroni-corrected pairwise comparisons where appropriate. Between-group effects (healthy vs. OSA) were evaluated separately for each physiological measure and sleep stage using Kruskal-Wallis and Wilcoxon tests. All analyses were performed in R (packages rstatix and tidyverse). Detailed test statistics and *p*-values for within-group stage effects, pairwise stage comparisons, and between-group differences are provided in Supplementary Tables S1-S3.

## 3 Results

DFA (Fig. 3) revealed distinct short– and long-term scaling behaviors across cardiovascular signals. Blood pressure (BP) recordings obtained with the Portapres system showed strong short-term correlations, with *α*_1_ ≈ 1.4 for both systolic and diastolic components, consistent with the behavior of RR intervals. A clear crossover from *α*_1_ *>* 1 to *α*_2_ *<* 1 was present, indicating strong local coupling and reduced short-term variability. In contrast, pulse arrival times (PAT) exhibited markedly lower short-term exponents and relatively stronger long-term correlations, resulting in a substantially reduced or absent crossover. This pattern suggests that PAT reflects a different temporal organization from pressure-derived measures, dominated by peripheral vascular influences rather than central arterial dynamics. The sleep-stage dependence of these differences for healthy subjects is summarized in Fig. 4.

**Fig 4.**
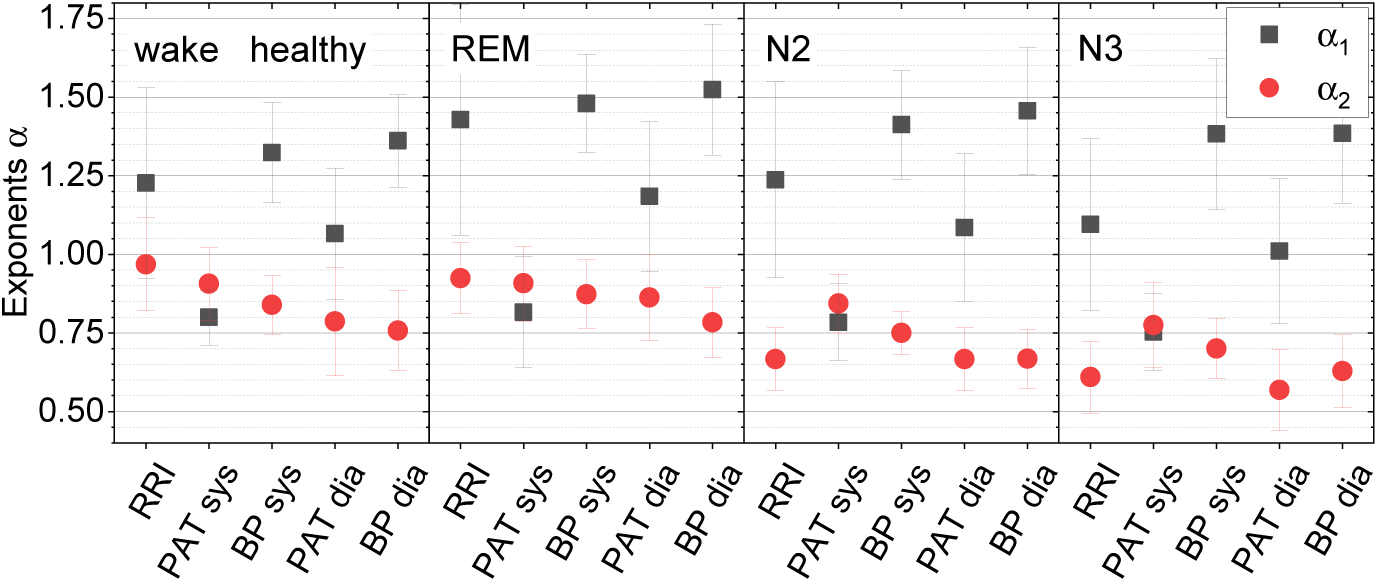
Short– and long-term scaling exponents for Portapres-derived measures in healthy subjects across sleep stages. Average short-term (*α*_1_, black) and long-term (*α*_2_, red) DFA exponents for RR intervals (RRI), systolic PAT (PAT_sys_), systolic BP (SBP), diastolic PAT (PAT_dia_), and diastolic BP (DBP) during wake, REM, light sleep (N2), and deep sleep (N3). Error bars denote standard deviations. For RRI and both systolic and diastolic BP, *α*_1_ was consistently and significantly larger than *α*_2_ across sleep stages, indicating a clear short– to long-term crossover. In contrast, PAT signals showed only weak or absent *α*_1_-*α*_2_ separation, with significant differences occurring only sporadically (primarily for PAT_dia_ in deep sleep), reflecting their different temporal organization compared with BP-derived measures.

Figure 4 shows the scaling exponents across sleep stages for the healthy group. RR intervals and both systolic and diastolic BP consistently exhibited a clear short– to long-term crossover, with *α*_1_ *>* 1 and *α*_2_ *<* 1 across all stages. This pattern reflects strong short-term cardiovascular coupling and reduced long-term persistence. In contrast, PAT_sys_ and PAT_dia_ showed only weak or absent separation between *α*_1_ and *α*_2_, with the diastolic PAT exhibiting a modest stage dependence only in deep sleep. These results confirm that, in healthy subjects, PAT captures substantially less short-term correlation structure than directly measured BP, consistent with the distinct temporal organization of PTT/PAT reported in earlier work [16].

Short– and long-term scaling patterns in the OSA group are shown in Fig. 5. As in healthy subjects, systolic and diastolic BP displayed a clear separation between *α*_1_ and *α*_2_ across sleep stages, while PAT signals again showed only weak or absent crossover. However, the magnitude of the *α*_1_-*α*_2_ difference was slightly reduced in OSA, particularly for RRI and for both systolic and diastolic BP during N2 and N3 sleep, consistent with a dampening of short-term cardiovascular coupling in the presence of sleep-disordered breathing.

**Fig 5.**
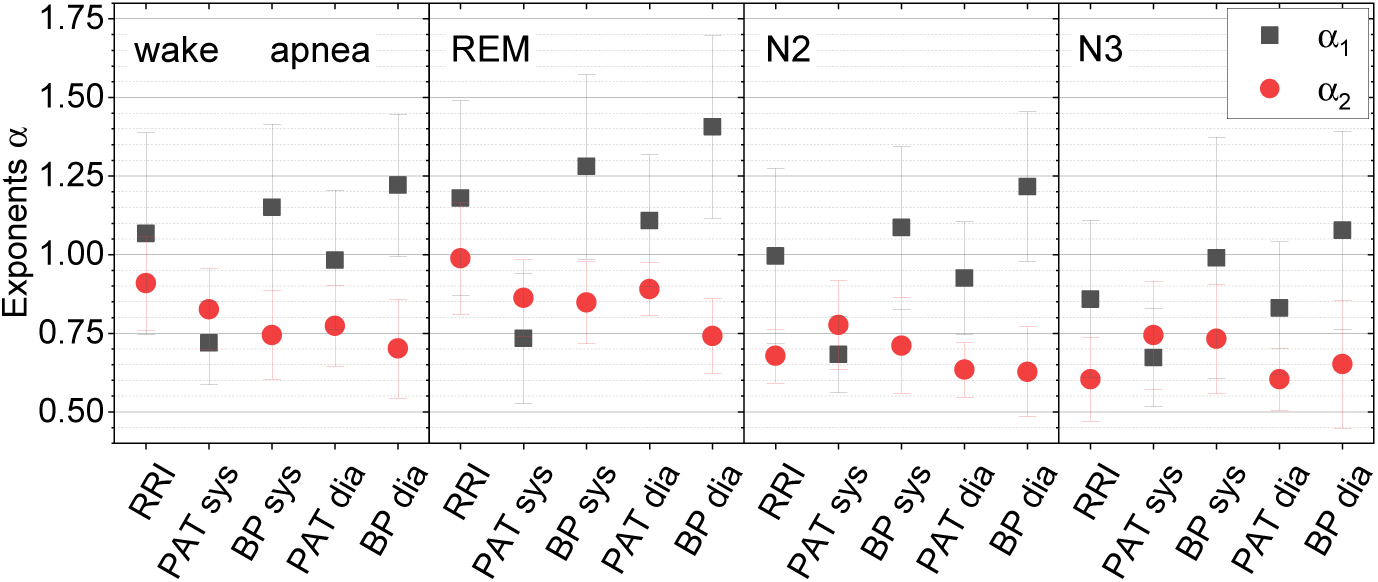
Short– and long-term scaling exponents for Portapres-derived measures in OSA subjects across sleep stages. Average short-term (*α*_1_, black) and long-term (*α*_2_, red) DFA exponents for RR intervals (RRI), systolic PAT (PAT_sys_), systolic BP (SBP), diastolic PAT (PAT_dia_), and diastolic BP (DBP) during wake, REM, light sleep (N2), and deep sleep (N3). Error bars denote standard deviations. As in healthy subjects, systolic and diastolic BP showed a clear separation between *α*_1_ and *α*_2_ across stages, whereas PAT signals again exhibited only weak or absent short– to long-term crossover, indicating that the altered scaling organization of PAT is preserved in OSA.

Direct comparison of the two groups confirmed these patterns. Systolic BP exhibited the most robust group differences, with consistently larger *α*_1_ and lower *α*_2_ in healthy subjects across sleep stages. Diastolic BP showed a similar trend, though the *α*_1_-*α*_2_ separation was most pronounced in N3 sleep. PAT measures provide the weakest differentiation between healthy and OSA subjects: systolic PAT displayed minimal separation of *α*_1_ and *α*_2_ in both groups, while diastolic PAT showed modest group differences only in deep sleep.

Figure 6 depicts the corresponding scaling behavior for the PPG-derived signals. In healthy subjects during sleep, systolic PPG (PPG_sys_) exhibited a clear short– to long-term crossover (*α*_1_ *> α*_2_), with diastolic PPG showing a similar but slightly weaker pattern. In contrast, PPG-derived PAT measures displayed only modest or absent separation between *α*_1_ and *α*_2_, mirroring the reduced crossover observed for ECG-anchored PAT in the Portapres signals.

**Fig 6.**
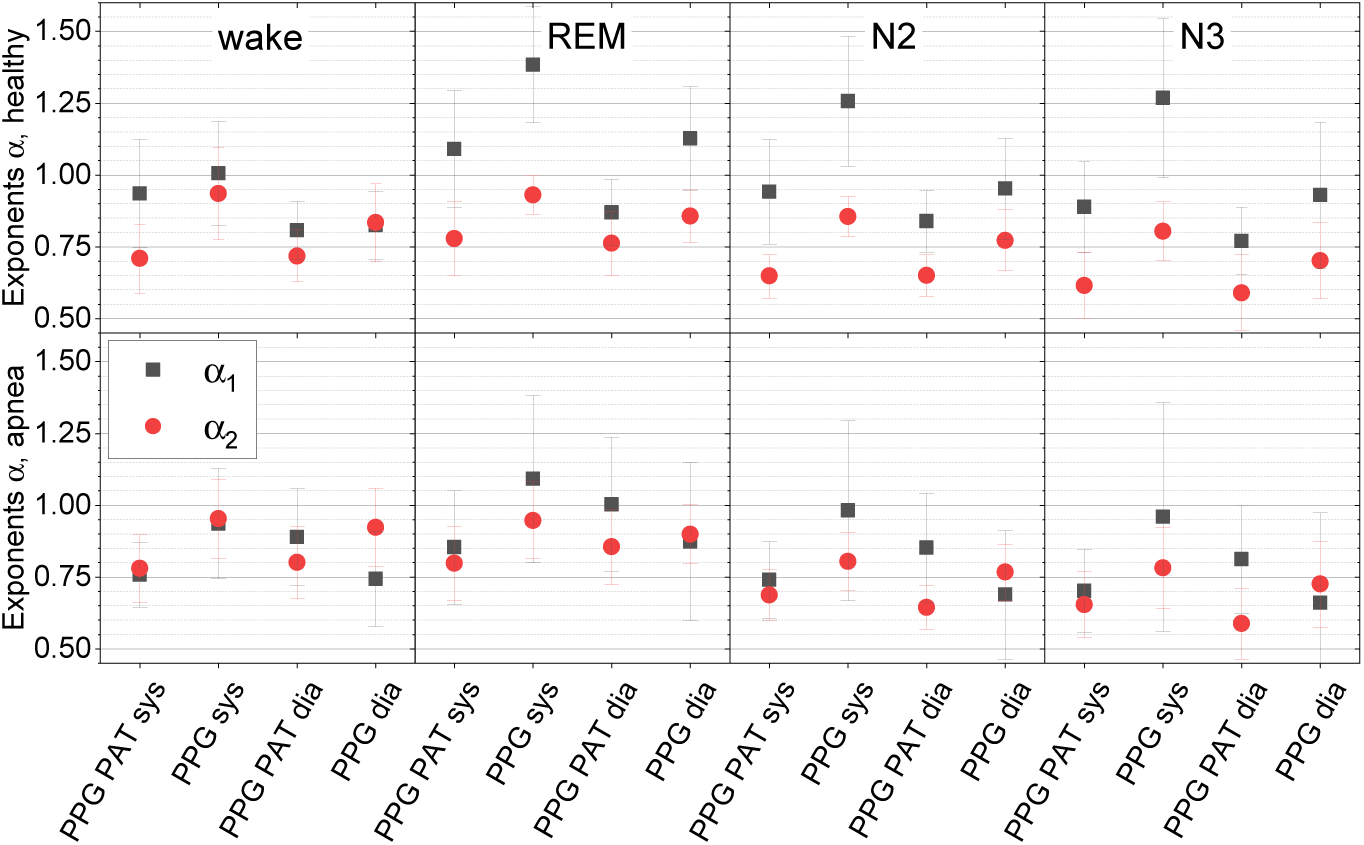
Short– and long-term scaling exponents for PPG-derived measures in healthy and OSA subjects. Average short-term (*α*_1_, black) and long-term (*α*_2_, red) DFA exponents for systolic PPG (PPG_sys_), diastolic PPG (PPG_dia_), systolic PAT (PPG-PAT_sys_), and diastolic PAT (PPG-PAT_dia_). Healthy participants (top row) showed a distinct *α*_1_-*α*_2_ crossover for PPG_sys_ and, to a lesser extent, PPG_dia_, whereas PAT-based indices exhibited much weaker separation. In OSA patients (bottom row), the *α*_1_-*α*_2_ crossover was substantially reduced or absent across all PPG-derived signals, indicating a loss of multiscale organization of peripheral pulse dynamics in the presence of sleep-disordered breathing. Error bars denote standard deviations.

In OSA patients, the temporal organization of the peripheral pulse wave was markedly changed: the *α*_1_-*α*_2_ crossover evident in healthy subjects was greatly diminished for both systolic and diastolic PPG, indicating a breakdown of short-term correlation structure in the presence of recurrent apneic events. PAT measures remained weakly organized in both groups, with only minor stage-dependent variations.

Together, the PPG results corroborate the Portapres findings and reinforce that PAT, whether derived from BP or PPG, does not reproduce the characteristic scaling properties of beat-to-beat BP fluctuations.

All qualitative patterns presented in this section were confirmed by non-parametric statistical testing; the full set of stage-wise and between-group comparisons is reported in the Supplementary Tables S1-S3.

## 4 Discussion

In this study, we examined the multiscale organization of cardiovascular dynamics across sleep stages by comparing DFA scaling exponents of BP-, PPG-, and PAT-derived measures in healthy individuals and patients with OSA. Our analysis addressed the central question raised by earlier work [16]: whether PAT/PTT-type timing measures reproduce the characteristic scaling structure of beat-to-beat blood pressure fluctuations. The present results demonstrate that they do not. Across both modalities (Portapress and PPG) and subject groups (healthy vs. OSA), only direct blood pressure signals as measured by Portapress exhibited a robust and physiologically meaningful short– to long-term crossover. PAT-based indices showed substantially weaker short-term correlations and reduced stage-dependent structure, highlighting their limited utility as surrogates for BP in fractal or scaling-based analyses.

### PAT does not replicate BP scaling: signal– and group-level interpretation

Across all sleep stages, both systolic and diastolic blood pressure (BP_sys_ and BP_dia_) consistently showed a clear separation between short-term (*α*_1_) and long-term (*α*_2_) scaling exponents. This pronounced crossover reflects well-organized cardiovascular regulation, in which baroreflex activity and vascular resistance tightly constrain short-term variability, while slower fluctuations reflect more independent autonomic and hemodynamic influences. Although the crossover was slightly more pronounced in non-REM sleep (N2 and N3), the overall magnitude of the *α*_1_-*α*_2_ difference was comparable for systolic and diastolic BP across all stages. This indicates that both systolic and diastolic pressure dynamics preserve the expected multiscale structure of beat-to-beat blood pressure regulation.

By contrast, the two PAT measures exhibited distinct behaviors. PAT_sys_ showed virtually no separation between *α*_1_ and *α*_2_ in any sleep stage, indicating an absence of meaningful short-term correlation structure. PAT_dia_ displayed a slightly clearer crossover, but this effect was modest and largely restricted to non-REM sleep. These findings mirror earlier observations [16] and highlight the mixed cardiovascular nature of PAT: because it includes part of the pre-ejection period (PEP), its short-term dynamics are influenced not only by vascular pulse-wave propagation, but also by cardiac contractility, autonomic modulation, and respiratory effects [11, 31]. As a result, PAT exhibits weaker and more variable short-term correlations than directly measured blood pressure, limiting its ability to reproduce the robust *α*_1_-*α*_2_ crossover characteristic of pressure-derived measures.

Between-group comparisons reinforced these differences. Patients with OSA exhibited a flattened scaling profile, with reduced *α*_1_ and a diminished short– to long-term crossover, particularly for BP_sys_. This pattern suggests impaired short-term autonomic adaptability and reduced baroreflex responsiveness during sleep. Diastolic BP showed similar but slightly weaker tendencies, whereas PAT-based indices again revealed only modest group differences. Overall, BP-derived scaling exponents remained the most effective discriminators between healthy and OSA subjects, while PAT indices showed limited diagnostic sensitivity.

PPG-derived measures displayed fundamentally different behavior from BP. In healthy subjects during sleep, PPG_sys_ exhibited a clear *α*_1_-*α*_2_ crossover, although weaker than that observed for pressure-derived signals. This partial retention of short-term structure indicates some correspondence to BP dynamics, likely mediated by shared autonomic inputs. In OSA, however, this crossover was largely absent, reflecting a loss of short-term organization in peripheral blood-volume dynamics. PAT measures derived from PPG showed the weakest and most inconsistent scaling structure, mirroring the behavior of ECG-referenced PAT derived from the Portapres signal.

Overall, these results demonstrate that PAT, regardless of whether it is derived from BP or PPG, does not reproduce the multiscale organization characteristic of beat-to-beat blood pressure fluctuations. PPG and PAT provide complementary information about peripheral vascular and autonomic function, but neither should be considered a surrogate for BP in fractal or scaling-based analyses.

### Measurement considerations

Continuous blood pressure in the DAPHNet dataset was recorded with the Portapres system, which inherently introduces characteristic calibration artifacts (Fig. 2). To prevent finger vasoconstriction, the device alternates the measured finger approximately every 30 minutes, often producing abrupt baseline offsets. In addition, the system performs periodic physiological calibrations every 10–70 beats, depending on signal quality, and occasional signal dropouts may occur [26, 27]. All affected segments were removed during preprocessing.

Because calibration shifts and finger switching disrupt absolute pressure baselines, only relative beat-to-beat variability of systolic and diastolic pressure could be analyzed reliably. While this precludes interpretation of absolute BP levels, the remaining variability is physiologically meaningful: fluctuations in SBP and DBP provide insight into autonomic regulation and have been shown to reflect cardiovascular dysregulation in OSA [28]. Thus, although constrained by device-specific limitations, the present analysis captures the temporal organization of pressure dynamics relevant to autonomic and vascular control.

### Device differences

The present study used finger PPG, whereas our earlier work [16] used wrist PPG. Finger and wrist waveforms differ substantially: finger PPG shows higher amplitude and clearer morphology, whereas wrist PPG exhibits weaker correlations with BP and greater motion sensitivity [32]. Wrist-worn devices have also been shown to slightly overestimate systolic and underestimate diastolic pressure, in part due to vertical displacement relative to heart level [33]. These differences underscore the importance of sensor location when interpreting PPG-derived timing indices.

### Physiological interpretation

Several physiological mechanisms explain why PAT differs fundamentally from BP in its scaling behavior. Many earlier studies referred to “PTT” while in fact analyzing ECG-referenced PAT (PTT^∗^), a distinction that is essential because the pre-ejection period (PEP) contributes substantially to PAT and introduces strong autonomic and cardiac influences [11, 31]. Geddes et al. [34] showed that PAT decreases with rising diastolic pressure, but with weaker correlation than true PTT, particularly at distal measurement sites. Payne et al. [31] further demonstrated that PEP can account for up to one-third of PAT variability, illustrating why PAT is an unreliable proxy for arterial stiffness or instantaneous blood pressure, and why its short-term fluctuations do not mirror those of peripheral pressure.

The PPG waveform additionally reflects local vascular compliance, wave-reflection properties, and sympathetic modulation [35]. The dicrotic notch, first characterized by Dillon [36], attenuates with arterial stiffening, altering waveform shape and thereby influencing scaling behavior. Respiratory and autonomic oscillations impose further low-frequency correlations in the waveform [37–39]. Vascular reflection parameters and stiffness indices also vary with smooth muscle tone, posture, and vasoactive modulation [40, 41]. Chen et al. [42] showed that arterial elasticity introduces variability in pulse-wave velocity that is independent of blood pressure, complicating interpretation of PTT/PAT-based estimates.

PAT itself is sensitive to autonomic modulation and posture. Recent work demonstrated that its short-term scaling exponent increases in the supine compared to the sitting position, reflecting greater autonomic stability when lying down [17]. This reinforces that PAT scaling reflects a combined influence of autonomic tone, cardiac electromechanical delay, and vascular propagation, not blood pressure alone.

Taken together, these mechanisms explain why PAT lacks the robust short-term scaling structure observed in BP and why PPG-derived indices differ fundamentally from pressure-derived measures. The observed scaling patterns reflect the interplay of cardiac delay, vascular propagation dynamics, and peripheral autonomic modulation, and therefore should not be interpreted as direct indicators of arterial stiffness or beat-to-beat blood pressure regulation.

## Conclusion

This study examined the multiscale organization of cardiovascular dynamics during sleep by comparing DFA-derived scaling exponents from continuous blood pressure (BP), PPG, and PAT signals in healthy individuals and patients with obstructive sleep apnea (OSA). Among all modalities, systolic and diastolic BP exhibited the clearest and most consistent separation between short– and long-term scaling exponents, reflecting robust autonomic and vascular regulation in healthy sleep and its attenuation in OSA. These findings demonstrate that beat-to-beat BP dynamics preserve a physiologically meaningful crossover structure that is highly sensitive to sleep-disordered breathing.

In contrast, PPG-derived signals showed a fundamentally different scaling organization. Although systolic PPG retained a partial *α*_1_-*α*_2_ crossover in healthy subjects, this structure was markedly weaker than in BP and largely absent in OSA, underscoring that PPG reflects peripheral blood-volume dynamics rather than central pressure regulation. PAT-based indices, whether derived from BP or from PPG, displayed the weakest and most inconsistent short-term correlations, largely due to their dependence on the pre-ejection period and other cardiac influences. As a result, PAT does not replicate BP scaling and should not be used as a surrogate for blood pressure in fractal or multiscale analyses.

In summary, our results highlight BP scaling as a sensitive marker of cardiovascular dysregulation in OSA, while PPG and PAT provide complementary but physiologically distinct information related to vascular tone, autonomic modulation, and cardiac electromechanical delay. Future work should explore multiscale coupling across combined cardiovascular and respiratory signals, and evaluate whether integrating BP-, PPG-, and PAT-based metrics can improve non-invasive screening strategies in sleep medicine.

## Acknowledgments

The study utilized previously collected data from [22], which was reanalyzed in accordance with the original ethical approval obtained from the study participants. We thank Dana Buck and Wibke Bartels for correspondence on their original research. KB gratefully acknowledges funding from the Minerva Foundation for supporting his research stay in Israel.

## Data availability

Data access is available from the authors upon reasonable request.

## Author contribution

KB performed the data preparation, preprocessing, initial analyses, and drafted the first version of the manuscript. JWK computed the DFA parameters and generated all DFA-related figures. MG contributed to the original data acquisition and provided methodological expertise on Portapres recordings and data preprocessing. TP supported the literature review as well as contextualization and physiological interpretation of the findings. NW conducted the statistical analyses, incl. all group comparisons and generated the p-value tables. KB, JWK, TP, and RPB jointly designed the study. RPB supervised the project as principal investigator and wrote the final draft of the manuscript. All authors discussed the results and contributed to manuscript revision, quality control, and final approval.

## Conflict of interest

The corresponding authors declare no conflict of interest.

## Supporting information

### Detailed signal preprocessing

All analyses were performed in MATLAB (R2023a, The MathWorks Inc., Natick, MA, USA). For each subject, signals were truncated to the time span covered by the Portapres blood pressure (BP) recording to ensure temporal alignment across modalities. The following channels were analyzed in detail: electrocardiogram (ECG, *f_s_* = 200 Hz), finger photoplethysmography (PPG, *f_s_* = 100 Hz), and continuous finger BP from the Portapres system (Model 2, FMS, The Netherlands; volume-clamp method, *f_s_* = 200 Hz). Where required, signals were resampled to a common time base using linear interpolation while preserving the original timing of R-peaks and pulse features.

#### ECG preprocessing and R-peak detection

ECG signals were first corrected for baseline wander. Baseline drift was estimated by applying a 0.25 s median filter to suppress QRS complexes, followed by a 0.75 s median filter to attenuate P and T waves. The resulting baseline estimate was then smoothed using a zero-phase, fifth-order Chebyshev Type II low-pass filter with a 4 Hz cutoff and subtracted from the original ECG. R-peaks were detected using the algorithm of Benítez et al. [24], and visually inspected for gross errors. The resulting R–R intervals (RRIs) served as the temporal reference for segmenting BP and PPG beats and for constructing the RRI time series used in DFA.

#### PPG preprocessing

PPG signals were preprocessed using a custom MATLAB routine designed to suppress high-frequency noise while preserving the physiological waveform. A fourth-order zero-phase Butterworth high-pass filter with a cutoff frequency of 10 Hz was applied to estimate the high-frequency noise component, which was then subtracted from the original signal. This procedure effectively reduced motion and sensor noise while maintaining the morphology of the pulsatile component.

#### BP preprocessing

For the Portapres BP signal, the same filtering procedure as for PPG was used, with the cutoff frequency of the high-pass filter set to 20 Hz and the sampling rate set to the native 200 Hz. This removed high-frequency artifacts while retaining the relevant physiological frequency range for systolic and diastolic pressure waveforms. In addition, calibration artifacts, finger-switching events, and brief signal dropouts were identified using the Portapres status flags and by visual inspection (see Fig. 2). Segments containing such artifacts were excluded from further analysis.

### Beat segmentation, feature extraction, and validation

Beat-level segmentation and validation were performed to ensure morphological consistency across both BP and PPG signals.

#### BP beats and fiducial points

For the BP signal, individual beats were defined using consecutive ECG R-peaks as temporal boundaries. Each beat was extended by one full R–R interval beyond the subsequent R-peak to include the late diastolic phase. The extracted segments were linearly resampled to 150 points to obtain phase-aligned waveforms across cardiac cycles.

For each subject, an ideal reference waveform was computed as the mean of all resampled beats. Morphological similarity between each individual beat and the reference waveform was quantified using the Pearson correlation coefficient. Beats with similarity ≥ 0.8 were considered morphologically valid and retained for further analysis. Systolic (SBP) and diastolic (DBP) points were initially identified by mapping the maximum and minimum of the reference waveform onto each original beat and then refined within a ±0.2 s window to ensure precise localization. Beats with SBP *>* 200 mmHg or DBP *<* 10 mmHg were excluded as physiologically implausible. The remaining beats were used to construct SBP and DBP time series.

#### PPG beats and fiducial points

PPG beats were segmented relative to ECG R-peaks to maintain synchrony between cardiac electrical activity and the peripheral pulse response. Consecutive R-peaks defined one complete PPG pulse cycle, and each segment was resampled to 200 points to standardize beat duration. For each subject, a reference PPG waveform was obtained by averaging all resampled beats. Morphological similarity between each beat and the reference was again quantified via the Pearson correlation coefficient; beats with similarity ≥ 0.6 were retained. Within each valid beat, the first local maximum after the onset was identified as the systolic PPG peak (PPG_sys_), and the subsequent local minimum on the falling limb was identified as the diastolic PPG minimum (PPG_dia_). Beats lacking a valid maximum–minimum pair or failing the similarity criterion were discarded. Segments affected by calibration, finger switching, or signal loss were also removed (Fig. 2).

#### Construction and validation of timing series

Pulse arrival times (PAT) were derived as described in the main Methods. Briefly, for Portapres BP signals:

- PAT_sys_: interval from the ECG R-peak to the SBP maximum,
- PAT_dia_: interval from the ECG R-peak to the diastolic BP minimum (DBP).

For PPG signals, PAT_sys_ and PAT_dia_ were defined analogously as the intervals from the R-peak to the PPG systolic peak and to the first diastolic PPG minimum following that peak, respectively. The following validation criteria were applied to the resulting beat-to-beat time series:

- **RRIs**: Interbeat intervals shorter than 0.33 s, longer than 2.0 s, 30% shorter than the previous RRI, or 60% longer than the previous RRI were discarded as non-normal beats, following previous work [16, 25].
- **Systolic PATs**: For Portapres-derived PAT_sys_, intervals shorter than 0.1 s, longer than 0.8 s, or 50% shorter/longer than the previous value were rejected. For PPG-derived PAT_sys_, the corresponding time limits were extended to −0.3 s and 1.0 s, reflecting somewhat less reliable overall signal timing.
- **Diastolic PATs**: For both Portapres– and PPG-derived PAT_dia_, intervals shorter than 0.3 s, longer than 3.0 s, or differing by more than ±50% from the previous value were excluded.
- **Blood pressure values**: BP values below 25 mmHg or above 250 mmHg were discarded as non-physiological.

After removing all invalid data points (beats), uninterrupted sequences with a minimum length of 20 consecutive measurements were retained for DFA.

### DFA implementation

Detrended fluctuation analysis (DFA), originally introduced by Peng et al. [29] and later extended to higher-order polynomial detrending [43], was used to quantify correlations in noisy, nonstationary time series across multiple time scales *s* [30]. The integrated signal of length *N* is divided into non-overlapping segments of size *s*. Within each segment, a polynomial trend is removed, and the mean-square residuals are averaged to obtain the fluctuation function *F* (*s*). For long-range correlated data, *F* (*s*) ∼ *s^α^*, with scaling exponent *α >* 0.5. This relation corresponds to a power-law in the power spectrum *P* (*f*) ∼ *f* ^−^*^β^*, where *β* = 2*α* − 1 [44]. In stationary cases (*α <* 1), this is equivalent to an autocorrelation function scaling *C*(*s*) ∼ *s*^−^*^γ^* with *γ* = 1 − *β* [45]. DFA is preferred over conventional spectral or autocorrelation methods because of its robustness to trends and nonstationarities [46, 47]. In the present analysis, DFA with second-order polynomial detrending (DFA2) was applied to all nine time series (RRIs; Portapres-derived SBP, DBP, PAT_sys_, PAT_dia_; and PPG-derived PPG_sys_, PPG_dia_, PAT_sys_, PAT_dia_) separately for each subject and sleep stage. Fluctuation functions were computed over beat-based window sizes and averaged across uninterrupted episodes using statistical weights proportional to episode duration. Short-term scaling exponents *α*_1_ were estimated over window sizes corresponding to 6–16 beats, and long-term exponents *α*_2_ over 50–200 beats, as in previous work [16]. Only fits with coefficient of determination *r*^2^ *>* 0.9 were retained. The resulting *α*_1_ and *α*_2_ values were then used in the group– and stage-level statistical analyses.

### Statistical analysis

Normality of *α*_1_ and *α*_2_ distributions was assessed using the Shapiro-Wilk test; because most distributions deviated significantly from normality (*p <* 0.05), non-parametric methods were used throughout. Within each subject group (healthy, OSA), Kruskal-Wallis tests were applied across the five Portapres-derived measures (RRI, SBP, DBP, PAT_sys_, PAT_dia_) and the four corresponding PPG-derived measures for each sleep stage. Significant effects were followed by Bonferroni-corrected pairwise Wilcoxon tests to identify specific stage contrasts. Stage-related differences within each physiological measure were assessed using Kruskal-Wallis tests across wake, REM, N2, and N3, again followed by Bonferroni-corrected pairwise Wilcoxon tests where appropriate. Between-group differences (healthy vs. OSA) were evaluated separately for each physiological measure and sleep stage using Kruskal-Wallis and Wilcoxon tests. All analyses were performed in R (packages rstatix and tidyverse). The complete statistical results are provided in Tables 2–4.

### Within-group stage effects

**Table 2.** Kruskal–Wallis tests across sleep stages within each group (DFA2). P-values are reported per signal and *α* exponent; *N* is the number of observations used.

| $\alpha$ | Signal | Group | KW $p$ | $N$ |
| --- | --- | --- | --- | --- |
| $\alpha_1$ | BP<sub>dia</sub> | Apnoea | 0.00669 | 72 |
| $\alpha_1$ | BP<sub>dia</sub> | Healthy | 0.04450 | 92 |
| $\alpha_1$ | PPG<sub>sys</sub> | Apnoea | 0.29600 | 72 |
| $\alpha_1$ | PPG<sub>sys</sub> | Healthy | 4.00e−06 | 92 |
| $\alpha_1$ | PPG<sub>dia</sub> | Apnoea | 0.03710 | 72 |
| $\alpha_1$ | PPG<sub>dia</sub> | Healthy | 1.59e−05 | 92 |
| $\alpha_1$ | PAT<sub>dia</sub> | Apnoea | 0.00261 | 72 |
| $\alpha_1$ | PAT<sub>dia</sub> | Healthy | 0.13000 | 92 |
| $\alpha_1$ | PPG-PAT<sub>sys</sub> | Apnoea | 0.12800 | 72 |
| $\alpha_1$ | PPG-PAT<sub>sys</sub> | Healthy | 0.00538 | 92 |
| $\alpha_1$ | PPG-PAT<sub>dia</sub> | Apnoea | 0.02230 | 72 |
| $\alpha_1$ | PPG-PAT<sub>dia</sub> | Healthy | 0.00877 | 92 |
| $\alpha_1$ | PAT<sub>sys</sub> | Apnoea | 0.80900 | 72 |
| $\alpha_1$ | PAT<sub>sys</sub> | Healthy | 0.69300 | 92 |
| $\alpha_1$ | BP<sub>sys</sub> | Apnoea | 0.04970 | 72 |
| $\alpha_1$ | BP<sub>sys</sub> | Healthy | 0.00868 | 92 |
| $\alpha_1$ | RRI | Apnoea | 0.02260 | 72 |
| $\alpha_1$ | RRI | Healthy | 0.01480 | 92 |
| $\alpha_2$ | BP<sub>dia</sub> | Apnoea | 0.13100 | 65 |
| $\alpha_2$ | BP<sub>dia</sub> | Healthy | 5.72e−05 | 91 |
| $\alpha_2$ | PPG<sub>sys</sub> | Apnoea | 1.41e−04 | 71 |
| $\alpha_2$ | PPG<sub>sys</sub> | Healthy | 1.07e−04 | 91 |
| $\alpha_2$ | PPG<sub>dia</sub> | Apnoea | 1.53e−05 | 71 |
| $\alpha_2$ | PPG<sub>dia</sub> | Healthy | 2.89e−04 | 91 |
| $\alpha_2$ | PAT<sub>dia</sub> | Apnoea | 9.86e−08 | 63 |
| $\alpha_2$ | PAT<sub>dia</sub> | Healthy | 1.53e−08 | 89 |
| $\alpha_2$ | PPG-PAT<sub>sys</sub> | Apnoea | 0.00209 | 68 |
| $\alpha_2$ | PPG-PAT<sub>sys</sub> | Healthy | 7.76e−04 | 81 |
| $\alpha_2$ | PPG-PAT<sub>dia</sub> | Apnoea | 1.98e−07 | 68 |
| $\alpha_2$ | PPG-PAT<sub>dia</sub> | Healthy | 1.25e−05 | 89 |
| $\alpha_2$ | PAT<sub>sys</sub> | Apnoea | 0.15900 | 63 |
| $\alpha_2$ | PAT<sub>sys</sub> | Healthy | 0.00302 | 89 |
| $\alpha_2$ | BP<sub>sys</sub> | Apnoea | 0.05500 | 65 |
| $\alpha_2$ | BP<sub>sys</sub> | Healthy | 4.90e−07 | 92 |
| $\alpha_2$ | RRI | Apnoea | 1.27e−09 | 69 |
| $\alpha_2$ | RRI | Healthy | 4.07e−13 | 92 |

### Pairwise stage comparisons

**Table 3.**
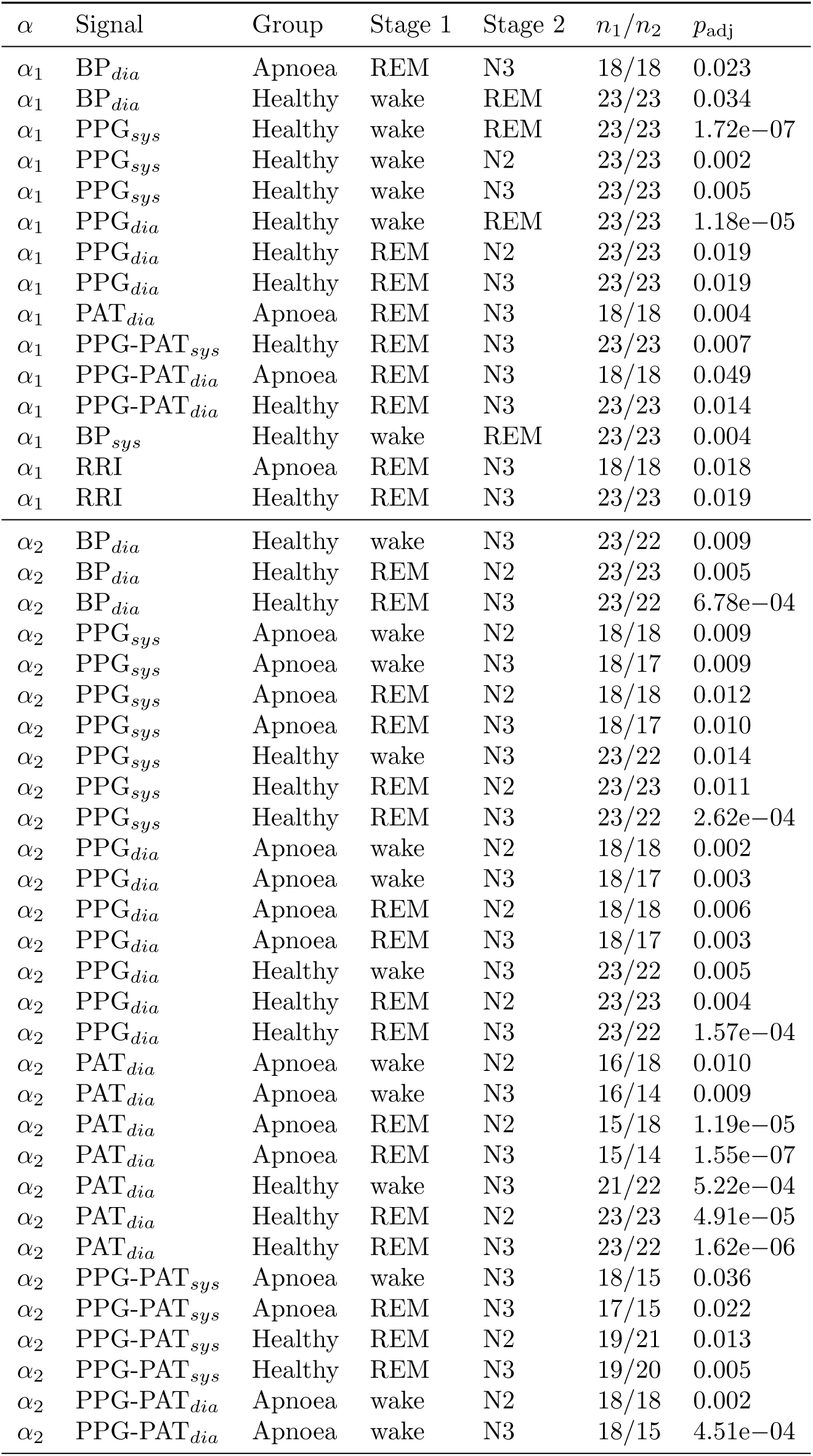
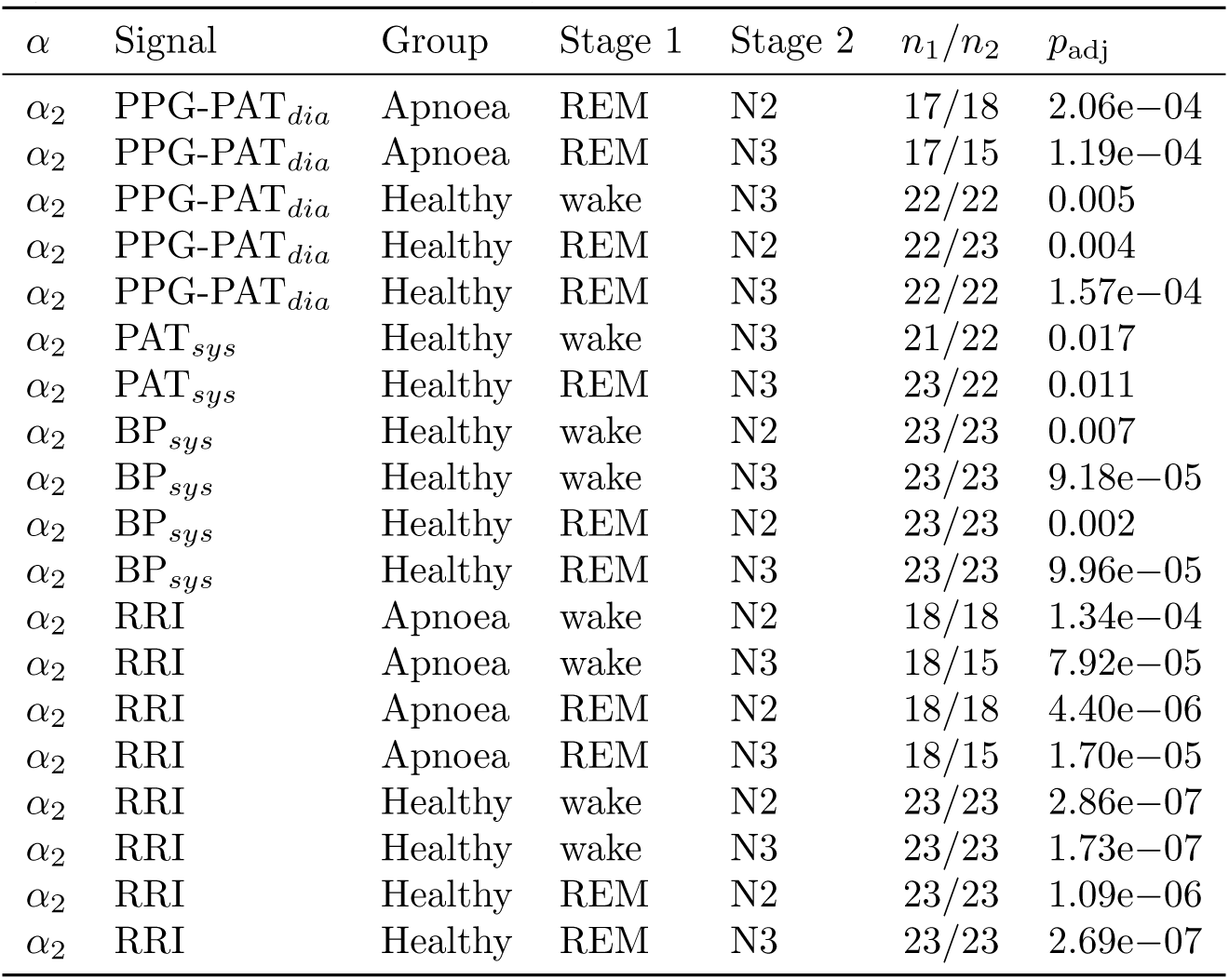
Significant Bonferroni-adjusted pairwise Wilcoxon tests across stages within groups (DFA2).

| $\alpha$ | Signal | Group | Stage 1 | Stage 2 | $n_1/n_2$ | $p_{\text{adj}}$ |
| --- | --- | --- | --- | --- | --- | --- |
| $\alpha_1$ | BP<sub>dia</sub> | Apnoea | REM | N3 | 18/18 | 0.023 |
| $\alpha_1$ | BP<sub>dia</sub> | Healthy | wake | REM | 23/23 | 0.034 |
| $\alpha_1$ | PPG<sub>sys</sub> | Healthy | wake | REM | 23/23 | 1.72e-07 |
| $\alpha_1$ | PPG<sub>sys</sub> | Healthy | wake | N2 | 23/23 | 0.002 |
| $\alpha_1$ | PPG<sub>sys</sub> | Healthy | wake | N3 | 23/23 | 0.005 |
| $\alpha_1$ | PPG<sub>dia</sub> | Healthy | wake | REM | 23/23 | 1.18e-05 |
| $\alpha_1$ | PPG<sub>dia</sub> | Healthy | REM | N2 | 23/23 | 0.019 |
| $\alpha_1$ | PPG<sub>dia</sub> | Healthy | REM | N3 | 23/23 | 0.019 |
| $\alpha_1$ | PAT<sub>dia</sub> | Apnoea | REM | N3 | 18/18 | 0.004 |
| $\alpha_1$ | PPG-PAT<sub>sys</sub> | Healthy | REM | N3 | 23/23 | 0.007 |
| $\alpha_1$ | PPG-PAT<sub>dia</sub> | Apnoea | REM | N3 | 18/18 | 0.049 |
| $\alpha_1$ | PPG-PAT<sub>dia</sub> | Healthy | REM | N3 | 23/23 | 0.014 |
| $\alpha_1$ | BP<sub>sys</sub> | Healthy | wake | REM | 23/23 | 0.004 |
| $\alpha_1$ | RRI | Apnoea | REM | N3 | 18/18 | 0.018 |
| $\alpha_1$ | RRI | Healthy | REM | N3 | 23/23 | 0.019 |
| $\alpha_2$ | BP<sub>dia</sub> | Healthy | wake | N3 | 23/22 | 0.009 |
| $\alpha_2$ | BP<sub>dia</sub> | Healthy | REM | N2 | 23/23 | 0.005 |
| $\alpha_2$ | BP<sub>dia</sub> | Healthy | REM | N3 | 23/22 | 6.78e-04 |
| $\alpha_2$ | PPG<sub>sys</sub> | Apnoea | wake | N2 | 18/18 | 0.009 |
| $\alpha_2$ | PPG<sub>sys</sub> | Apnoea | wake | N3 | 18/17 | 0.009 |
| $\alpha_2$ | PPG<sub>sys</sub> | Apnoea | REM | N2 | 18/18 | 0.012 |
| $\alpha_2$ | PPG<sub>sys</sub> | Apnoea | REM | N3 | 18/17 | 0.010 |
| $\alpha_2$ | PPG<sub>sys</sub> | Healthy | wake | N3 | 23/22 | 0.014 |
| $\alpha_2$ | PPG<sub>sys</sub> | Healthy | REM | N2 | 23/23 | 0.011 |
| $\alpha_2$ | PPG<sub>sys</sub> | Healthy | REM | N3 | 23/22 | 2.62e-04 |
| $\alpha_2$ | PPG<sub>dia</sub> | Apnoea | wake | N2 | 18/18 | 0.002 |
| $\alpha_2$ | PPG<sub>dia</sub> | Apnoea | wake | N3 | 18/17 | 0.003 |
| $\alpha_2$ | PPG<sub>dia</sub> | Apnoea | REM | N2 | 18/18 | 0.006 |
| $\alpha_2$ | PPG<sub>dia</sub> | Apnoea | REM | N3 | 18/17 | 0.003 |
| $\alpha_2$ | PPG<sub>dia</sub> | Healthy | wake | N3 | 23/22 | 0.005 |
| $\alpha_2$ | PPG<sub>dia</sub> | Healthy | REM | N2 | 23/23 | 0.004 |
| $\alpha_2$ | PPG<sub>dia</sub> | Healthy | REM | N3 | 23/22 | 1.57e-04 |
| $\alpha_2$ | PAT<sub>dia</sub> | Apnoea | wake | N2 | 16/18 | 0.010 |
| $\alpha_2$ | PAT<sub>dia</sub> | Apnoea | wake | N3 | 16/14 | 0.009 |
| $\alpha_2$ | PAT<sub>dia</sub> | Apnoea | REM | N2 | 15/18 | 1.19e-05 |
| $\alpha_2$ | PAT<sub>dia</sub> | Apnoea | REM | N3 | 15/14 | 1.55e-07 |
| $\alpha_2$ | PAT<sub>dia</sub> | Healthy | wake | N3 | 21/22 | 5.22e-04 |
| $\alpha_2$ | PAT<sub>dia</sub> | Healthy | REM | N2 | 23/23 | 4.91e-05 |
| $\alpha_2$ | PAT<sub>dia</sub> | Healthy | REM | N3 | 23/22 | 1.62e-06 |
| $\alpha_2$ | PPG-PAT<sub>sys</sub> | Apnoea | wake | N3 | 18/15 | 0.036 |
| $\alpha_2$ | PPG-PAT<sub>sys</sub> | Apnoea | REM | N3 | 17/15 | 0.022 |
| $\alpha_2$ | PPG-PAT<sub>sys</sub> | Healthy | REM | N2 | 19/21 | 0.013 |
| $\alpha_2$ | PPG-PAT<sub>sys</sub> | Healthy | REM | N3 | 19/20 | 0.005 |
| $\alpha_2$ | PPG-PAT<sub>dia</sub> | Apnoea | wake | N2 | 18/18 | 0.002 |
| $\alpha_2$ | PPG-PAT<sub>dia</sub> | Apnoea | wake | N3 | 18/15 | 4.51e-04 |
| $\alpha_2$ | PPG-PAT<sub>dia</sub> | Apnoea | REM | N2 | 17/18 | 2.06e−04 |
| $\alpha_2$ | PPG-PAT<sub>dia</sub> | Apnoea | REM | N3 | 17/15 | 1.19e−04 |
| $\alpha_2$ | PPG-PAT<sub>dia</sub> | Healthy | wake | N3 | 22/22 | 0.005 |
| $\alpha_2$ | PPG-PAT<sub>dia</sub> | Healthy | REM | N2 | 22/23 | 0.004 |
| $\alpha_2$ | PPG-PAT<sub>dia</sub> | Healthy | REM | N3 | 22/22 | 1.57e−04 |
| $\alpha_2$ | PAT<sub>sys</sub> | Healthy | wake | N3 | 21/22 | 0.017 |
| $\alpha_2$ | PAT<sub>sys</sub> | Healthy | REM | N3 | 23/22 | 0.011 |
| $\alpha_2$ | BP<sub>sys</sub> | Healthy | wake | N2 | 23/23 | 0.007 |
| $\alpha_2$ | BP<sub>sys</sub> | Healthy | wake | N3 | 23/23 | 9.18e−05 |
| $\alpha_2$ | BP<sub>sys</sub> | Healthy | REM | N2 | 23/23 | 0.002 |
| $\alpha_2$ | BP<sub>sys</sub> | Healthy | REM | N3 | 23/23 | 9.96e−05 |
| $\alpha_2$ | RRI | Apnoea | wake | N2 | 18/18 | 1.34e−04 |
| $\alpha_2$ | RRI | Apnoea | wake | N3 | 18/15 | 7.92e−05 |
| $\alpha_2$ | RRI | Apnoea | REM | N2 | 18/18 | 4.40e−06 |
| $\alpha_2$ | RRI | Apnoea | REM | N3 | 18/15 | 1.70e−05 |
| $\alpha_2$ | RRI | Healthy | wake | N2 | 23/23 | 2.86e−07 |
| $\alpha_2$ | RRI | Healthy | wake | N3 | 23/23 | 1.73e−07 |
| $\alpha_2$ | RRI | Healthy | REM | N2 | 23/23 | 1.09e−06 |
| $\alpha_2$ | RRI | Healthy | REM | N3 | 23/23 | 2.69e−07 |

### Between-group differences by stage

**Table 4.**
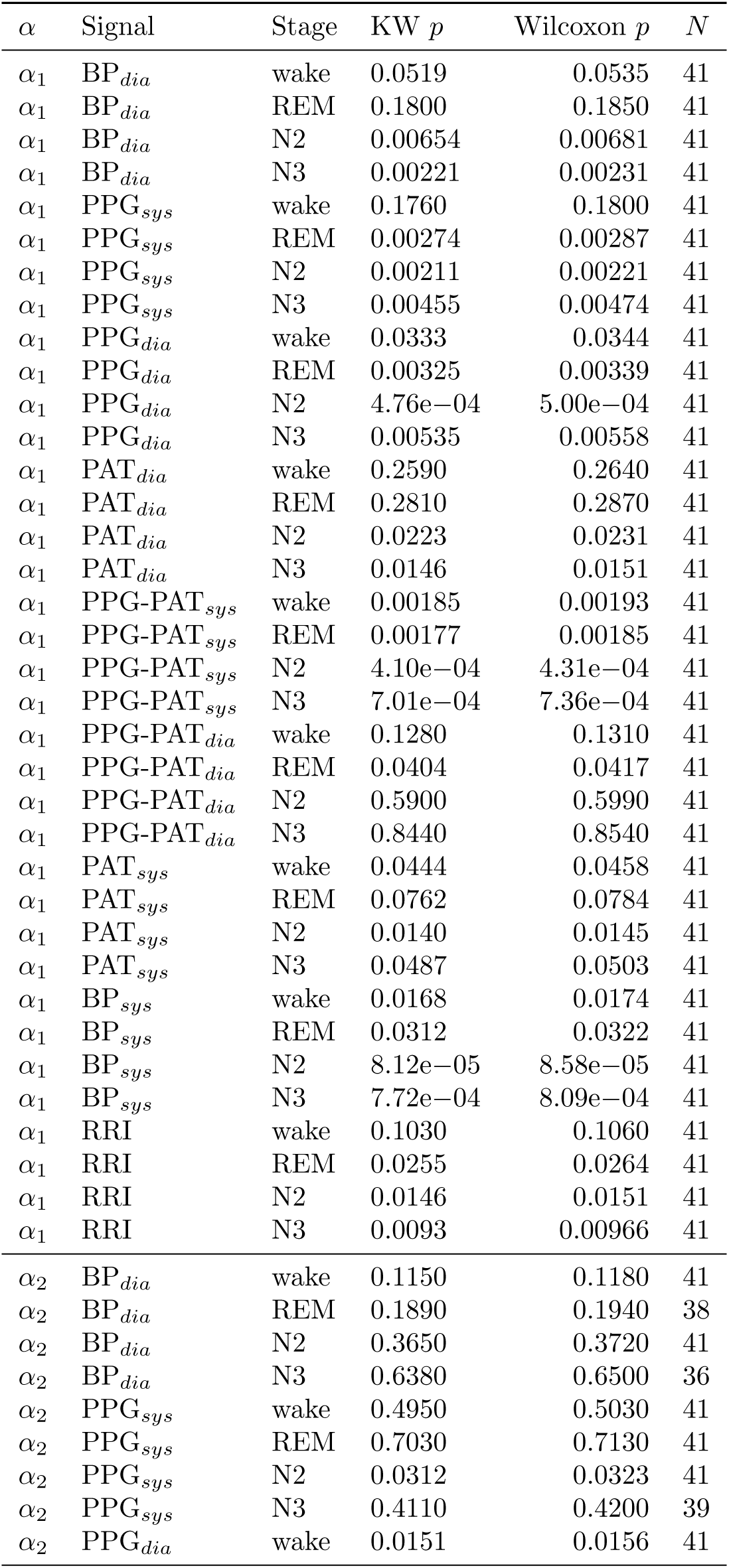
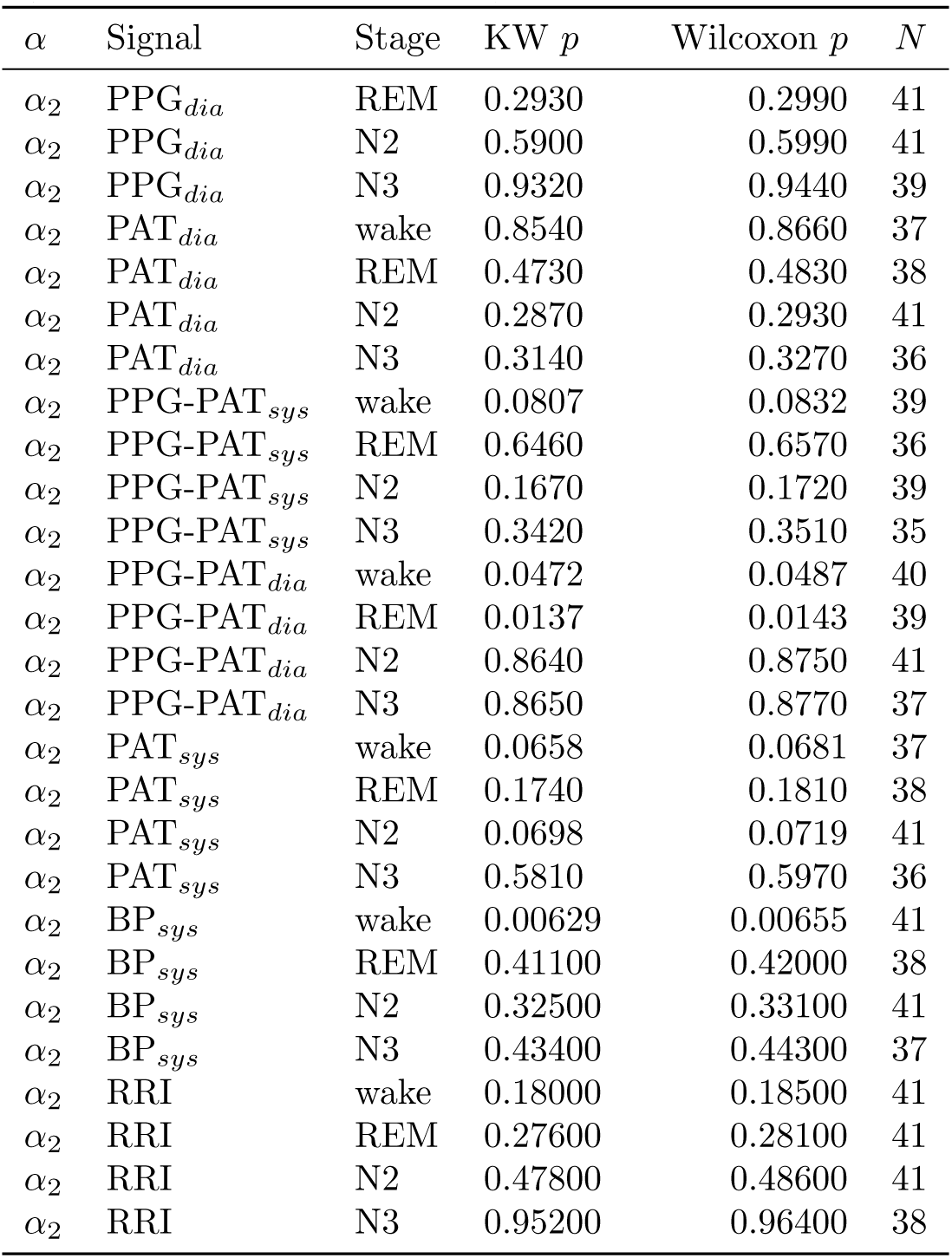
Between-group (Healthy vs. Apnea) comparisons per stage (DFA2). Both Kruskal-Wallis (KW) and Wilcoxon *p*-values are shown; *N* denotes total observations.

| $\alpha$ | Signal | Stage | KW $p$ | Wilcoxon $p$ | $N$ |
| --- | --- | --- | --- | --- | --- |
| $\alpha_1$ | BP<sub>dia</sub> | wake | 0.0519 | 0.0535 | 41 |
| $\alpha_1$ | BP<sub>dia</sub> | REM | 0.1800 | 0.1850 | 41 |
| $\alpha_1$ | BP<sub>dia</sub> | N2 | 0.00654 | 0.00681 | 41 |
| $\alpha_1$ | BP<sub>dia</sub> | N3 | 0.00221 | 0.00231 | 41 |
| $\alpha_1$ | PPG<sub>sys</sub> | wake | 0.1760 | 0.1800 | 41 |
| $\alpha_1$ | PPG<sub>sys</sub> | REM | 0.00274 | 0.00287 | 41 |
| $\alpha_1$ | PPG<sub>sys</sub> | N2 | 0.00211 | 0.00221 | 41 |
| $\alpha_1$ | PPG<sub>sys</sub> | N3 | 0.00455 | 0.00474 | 41 |
| $\alpha_1$ | PPG<sub>dia</sub> | wake | 0.0333 | 0.0344 | 41 |
| $\alpha_1$ | PPG<sub>dia</sub> | REM | 0.00325 | 0.00339 | 41 |
| $\alpha_1$ | PPG<sub>dia</sub> | N2 | 4.76e−04 | 5.00e−04 | 41 |
| $\alpha_1$ | PPG<sub>dia</sub> | N3 | 0.00535 | 0.00558 | 41 |
| $\alpha_1$ | PAT<sub>dia</sub> | wake | 0.2590 | 0.2640 | 41 |
| $\alpha_1$ | PAT<sub>dia</sub> | REM | 0.2810 | 0.2870 | 41 |
| $\alpha_1$ | PAT<sub>dia</sub> | N2 | 0.0223 | 0.0231 | 41 |
| $\alpha_1$ | PAT<sub>dia</sub> | N3 | 0.0146 | 0.0151 | 41 |
| $\alpha_1$ | PPG-PAT<sub>sys</sub> | wake | 0.00185 | 0.00193 | 41 |
| $\alpha_1$ | PPG-PAT<sub>sys</sub> | REM | 0.00177 | 0.00185 | 41 |
| $\alpha_1$ | PPG-PAT<sub>sys</sub> | N2 | 4.10e−04 | 4.31e−04 | 41 |
| $\alpha_1$ | PPG-PAT<sub>sys</sub> | N3 | 7.01e−04 | 7.36e−04 | 41 |
| $\alpha_1$ | PPG-PAT<sub>dia</sub> | wake | 0.1280 | 0.1310 | 41 |
| $\alpha_1$ | PPG-PAT<sub>dia</sub> | REM | 0.0404 | 0.0417 | 41 |
| $\alpha_1$ | PPG-PAT<sub>dia</sub> | N2 | 0.5900 | 0.5990 | 41 |
| $\alpha_1$ | PPG-PAT<sub>dia</sub> | N3 | 0.8440 | 0.8540 | 41 |
| $\alpha_1$ | PAT<sub>sys</sub> | wake | 0.0444 | 0.0458 | 41 |
| $\alpha_1$ | PAT<sub>sys</sub> | REM | 0.0762 | 0.0784 | 41 |
| $\alpha_1$ | PAT<sub>sys</sub> | N2 | 0.0140 | 0.0145 | 41 |
| $\alpha_1$ | PAT<sub>sys</sub> | N3 | 0.0487 | 0.0503 | 41 |
| $\alpha_1$ | BP<sub>sys</sub> | wake | 0.0168 | 0.0174 | 41 |
| $\alpha_1$ | BP<sub>sys</sub> | REM | 0.0312 | 0.0322 | 41 |
| $\alpha_1$ | BP<sub>sys</sub> | N2 | 8.12e−05 | 8.58e−05 | 41 |
| $\alpha_1$ | BP<sub>sys</sub> | N3 | 7.72e−04 | 8.09e−04 | 41 |
| $\alpha_1$ | RRI | wake | 0.1030 | 0.1060 | 41 |
| $\alpha_1$ | RRI | REM | 0.0255 | 0.0264 | 41 |
| $\alpha_1$ | RRI | N2 | 0.0146 | 0.0151 | 41 |
| $\alpha_1$ | RRI | N3 | 0.0093 | 0.00966 | 41 |
| $\alpha_2$ | BP<sub>dia</sub> | wake | 0.1150 | 0.1180 | 41 |
| $\alpha_2$ | BP<sub>dia</sub> | REM | 0.1890 | 0.1940 | 38 |
| $\alpha_2$ | BP<sub>dia</sub> | N2 | 0.3650 | 0.3720 | 41 |
| $\alpha_2$ | BP<sub>dia</sub> | N3 | 0.6380 | 0.6500 | 36 |
| $\alpha_2$ | PPG<sub>sys</sub> | wake | 0.4950 | 0.5030 | 41 |
| $\alpha_2$ | PPG<sub>sys</sub> | REM | 0.7030 | 0.7130 | 41 |
| $\alpha_2$ | PPG<sub>sys</sub> | N2 | 0.0312 | 0.0323 | 41 |
| $\alpha_2$ | PPG<sub>sys</sub> | N3 | 0.4110 | 0.4200 | 39 |
| $\alpha_2$ | PPG<sub>dia</sub> | wake | 0.0151 | 0.0156 | 41 |
| $\alpha_2$ | PPG<sub>dia</sub> | REM | 0.2930 | 0.2990 | 41 |
| $\alpha_2$ | PPG<sub>dia</sub> | N2 | 0.5900 | 0.5990 | 41 |
| $\alpha_2$ | PPG<sub>dia</sub> | N3 | 0.9320 | 0.9440 | 39 |
| $\alpha_2$ | PAT<sub>dia</sub> | wake | 0.8540 | 0.8660 | 37 |
| $\alpha_2$ | PAT<sub>dia</sub> | REM | 0.4730 | 0.4830 | 38 |
| $\alpha_2$ | PAT<sub>dia</sub> | N2 | 0.2870 | 0.2930 | 41 |
| $\alpha_2$ | PAT<sub>dia</sub> | N3 | 0.3140 | 0.3270 | 36 |
| $\alpha_2$ | PPG-PAT<sub>sys</sub> | wake | 0.0807 | 0.0832 | 39 |
| $\alpha_2$ | PPG-PAT<sub>sys</sub> | REM | 0.6460 | 0.6570 | 36 |
| $\alpha_2$ | PPG-PAT<sub>sys</sub> | N2 | 0.1670 | 0.1720 | 39 |
| $\alpha_2$ | PPG-PAT<sub>sys</sub> | N3 | 0.3420 | 0.3510 | 35 |
| $\alpha_2$ | PPG-PAT<sub>dia</sub> | wake | 0.0472 | 0.0487 | 40 |
| $\alpha_2$ | PPG-PAT<sub>dia</sub> | REM | 0.0137 | 0.0143 | 39 |
| $\alpha_2$ | PPG-PAT<sub>dia</sub> | N2 | 0.8640 | 0.8750 | 41 |
| $\alpha_2$ | PPG-PAT<sub>dia</sub> | N3 | 0.8650 | 0.8770 | 37 |
| $\alpha_2$ | PAT<sub>sys</sub> | wake | 0.0658 | 0.0681 | 37 |
| $\alpha_2$ | PAT<sub>sys</sub> | REM | 0.1740 | 0.1810 | 38 |
| $\alpha_2$ | PAT<sub>sys</sub> | N2 | 0.0698 | 0.0719 | 41 |
| $\alpha_2$ | PAT<sub>sys</sub> | N3 | 0.5810 | 0.5970 | 36 |
| $\alpha_2$ | BP<sub>sys</sub> | wake | 0.00629 | 0.00655 | 41 |
| $\alpha_2$ | BP<sub>sys</sub> | REM | 0.41100 | 0.42000 | 38 |
| $\alpha_2$ | BP<sub>sys</sub> | N2 | 0.32500 | 0.33100 | 41 |
| $\alpha_2$ | BP<sub>sys</sub> | N3 | 0.43400 | 0.44300 | 37 |
| $\alpha_2$ | RRI | wake | 0.18000 | 0.18500 | 41 |
| $\alpha_2$ | RRI | REM | 0.27600 | 0.28100 | 41 |
| $\alpha_2$ | RRI | N2 | 0.47800 | 0.48600 | 41 |
| $\alpha_2$ | RRI | N3 | 0.95200 | 0.96400 | 38 |

## Notes

### Competing Interest Statement

The authors have declared no competing interest.

